# *In vivo* electrophysiological validation of DREADD-based inhibition of pallidal neurons in the non-human primate

**DOI:** 10.1101/2020.01.03.893610

**Authors:** Marc Deffains, Tho Haï Nguyen, Hugues Orignac, Nathalie Biendon, Sandra Dovero, Erwan Bezard, Thomas Boraud

## Abstract

Designer Receptors Exclusively Activated by Designer Drugs (DREADDs) are widely used in rodents to manipulate neuronal activity and establish causal links between structure and function. Their utilization in non-human primates (NHPs) is however limited and their efficacy still debated. Here, we tested DREADD expression in the NHP external globus pallidus (GPe) and electrophysiologically validated DREADD-based inhibition of GPe neurons in the anesthetized monkey.To do so, we performed intracerebral injections of viral construct expressing hM4Di receptor under a neuron-specific promoter into the GPe. Then, we recorded the neuronal activity in the DREADD-transduced (test condition) and DREADD-free (control condition) GPe of two anesthetized animals following local intra-GPe microinjection of clozapine-N-oxide (CNO). In total, 19 and 8 well-isolated and stable units were recorded in the DREADD-transduced and DREADD-free GPe, respectively. Overall, we found that almost half (9/19) of the units modulated their activity following CNO injection in DREADD-transduced GPe. Surprisingly, neuronal activity of the GPe units exhibited diverse patterns in timing and polarity (increase/decrease) of firing rate modulations during and after CNO injection. Nevertheless, decreases were exclusive and stronger after CNO injection. In contrast, only one unit modulated its activity after CNO injection in DREADD-free GPe. Moreover, post-mortem histochemical analysis revealed that hM4Di DREADDs were expressed at high level in the GPe neurons located in the vicinity of the viral construct injection sites. Our results therefore show *in vivo* DREADD-based inhibition of pallidal neurons in the NHP model and reinforce the view that DREADD technology can be effective in NHPs.

## Introduction

Designer Receptors Exclusively Activated by Designer Drugs (DREADDs) technique can be used as a powerful tool to remotely, reversibly and repeatedly manipulate the neural mechanisms underlying higher brain functions with minimal invasiveness and maximal control of anatomical coverage (Roth, 2016).The hM4Di receptor (i.e., engineered version of the M4 muscarinic acetylcholine receptor) is a popular muscarinic-based DREADD that can reduce the activity of DREADD-transduced neurons when activated by clozapine-n-oxide (CNO). This DREADD technology is widely used in rodents (e.g., Ferguson *et al.*, 2011, 2013; Chang *et al.*, 2015; Smith *et al.*, 2016); however, its efficacy in non-human primates (NHPs) is still debated. CNO has low blood-brain permeability (Gomez *et al.*, 2017; Raper *et al.*, 2017) and is reverse-metabolized to clozapine (Jann *et al.*, 1994; Chang *et al.*, 1998; MacLaren *et al.*, 2016; Gomez *et al.*, 2017) which shows high blood-brain permeability and high affinity to muscarinic-based DREADDs (Ji *et al.*, 2016; Gomez *et al.*, 2017). However, unlike CNO that is considered an inert drug (without other known receptor interactions), clozapine is capable of binding to a variety of endogenous receptors (Galvan *et al.*, 2018). Moreover, a recent NHP study demonstrated that the level of plasma membrane expression of the DREADDs and consequently the efficacy of the DREADD approach depend on the nature of the protein tag fused to the DREADD (Galvan *et al.*, 2019). In NHP experiments the number of animals used is typically low and the possibility of repeated experiments to validate and/or adjust the technology is limited. This is probably why only few NHP studies using DREADDs have been conducted so far. (Eldridge *et al.*, 2016; Grayson *et al.*, 2016; Upright *et al.*, 2018; Bonaventura *et al.*, 2019; Galvan *et al.*, 2019). In fact, some NHP studies have already observed DREADD-based behavioral effects (Eldridge *et al.*, 2016; Grayson *et al.*, 2016; Upright *et al.*, 2018); however, there is still no direct evidence that neuronal activity can be manipulated in monkeys using DREADDs.

To fill this gap, we tested DREADD expression in the NHP external globus pallidus (GPe) and validated DREADD-based inhibition of GPe neurons using electrophysiological recordings in the anesthetized monkey. To do so, we transduced NHP GPe neurons with hM4Di receptor and recorded single-unit activity in the DREADD-transduced and DREADD-free (Controls) GPe following local intra-GPe microinjection in the proximity of the electrode tip (within ~2mm radius) of CNO.

## Materials and Methods

### Animals

Two rhesus monkeys (F and E, *Macaca mulatta*) weighing ~ 7-8kg, were used in this study. The monkeys were housed in the animal facility of the Institute of Neurodegenerative Diseases (UMR CNRS 5293, Bordeaux, France) under standard conditions (12/12h day/night cycle, ~ 60% humidity and ~ 22°C). All experimental protocols comply with Council Directive of 2010 (2010/63/EU) of the European Community and the National Institute of Health Guide for the Care and Use of Laboratory Animals. The proposed research has received agreement from French Ethical Committee for Animal Research CE50 (agreement number: C33063268).

### Viral constructs and surgical procedures

To transduce GPe neurons with hM4Di DREADDs, we used adeno-associated virus (AAV) expressing hM4Di receptor fused to mCherry fluorescent protein tag under human synapsin promoter [AAV2-hSyn-hM4D(Gi)-mCherry (3*10^12^ and 1.1*10^13^ viral particles/ml for monkeys F and E, respectively), Addgene, Cambridge, MA, USA].

Surgeries were performed under aseptic conditions and isoflurane and O2 deep anesthesia, following induction with ketamine (10mg/kg IM) and atropine (0.05mg/kg IM). Body temperature, heart rate blood pressure and SpO2 were monitored throughout all surgical procedures. During surgery, the head of the animal was fixed in a stereotaxic frame (M2e-Unimécanique, Eaubonne, Val-d’Oise, France) and craniotomies were made above GPe (left and right GPe in monkeys F and E, respectively). Viral construct were administered in 4 injections within GPe using a UMP3 microsyringe pump injector (World Precision Instruments, Sarasota, FL, USA) with 100µl Hamilton syringe. For each injection, a needle was lowered to the injection site and 20µl of the viral constructs were infused at a rate of 2 µl/min. Injection sites within GPe were ascertained according to the stereotaxic coordinates of GPe (Paxinos *et al.*, 2000) and guided by ventriculography and single-unit electrophysiological recordings. A ventriculographic cannula was introduced into the anterior horn of the lateral ventricle and a contrast medium (Omnipaque) was injected (Piron *et al.*, 2016). Corrections in the position of the injection sites were performed according to the line between the anterior and posterior commissures. Theoretical coordinates of the injection sites (in mm) from the vertical plane passing through the interaural line (AP), the midline (L) and the horizontal plane passing through the interaural line were: AP:18.3, L:5, D:16; AP:18.3, L:7, D:16; AP:16.5, L:6, D:16; AP:16.5, L:9, D:16 (Paxinos *et al.*, 2000). After each injection, there was a 5-min waiting period to minimize viral construct spread during retractions. Analgesics (Metacam, 0.2mg/kg IM) were administered during the first 3 postoperative days. A 4-week minimum waiting period for allowing DREADD expression was allotted before starting recordings.

### Electrophysiology

During the recording sessions, the animal was re-anesthetized (see above for the details) and its head was re-immobilized in the stereotaxic frame (M2e-Unimécanique, Eaubonne, Val-d’Oise, France). Narylene-coated tungsten microelectrodes (impedance at 1 kHz: ~1 MΩ) were manually advanced within the GPe to the viral construct injection sites. The electrical activity was amplified by 100, band-pass filtered from 250 to 6000Hz using a hardware two-pole Butterworth filter and sampled at 25kHz (advanced wireless W2100, Multichannel Systems, Reutlingen, BW, Germany)

After radiographic and electrophysiological confirmation of the electrode position, 25µl Hamilton syringe [mounted on a UMP3 microsyringe pump injector (World Precision Instruments, Sarasota, FL, USA)] was filled with CNO (Bio-Techne, Minneapolis, MN, USA) and needle was lowered to it final position (Fig.1A, B and C). When a neuron was found [within ~2mm radius from the tip of the needle, so that we could reasonably (Myers, 1966) conclude that the recorded neuron is located within the diffusion area of 5µl injected CNO], its activity was recorded during 10-min baseline period in order to assess the stability of its activity (before starting CNO injection). If its activity was sufficiently stable, 5µl CNO (at concentration of 1mM.) was injected at 0.2ul/min (25-min injection period) and recording stopped only 50min after starting CNO injection. To guarantee a complete CNO washout between the recordings, only one CNO injection was performed per day of recording. As controls, similar recordings following CNO injections were made in DREADD-free GPe (i.e., right and left GPe in monkeys F and E, respectively).

**Fig.1.**
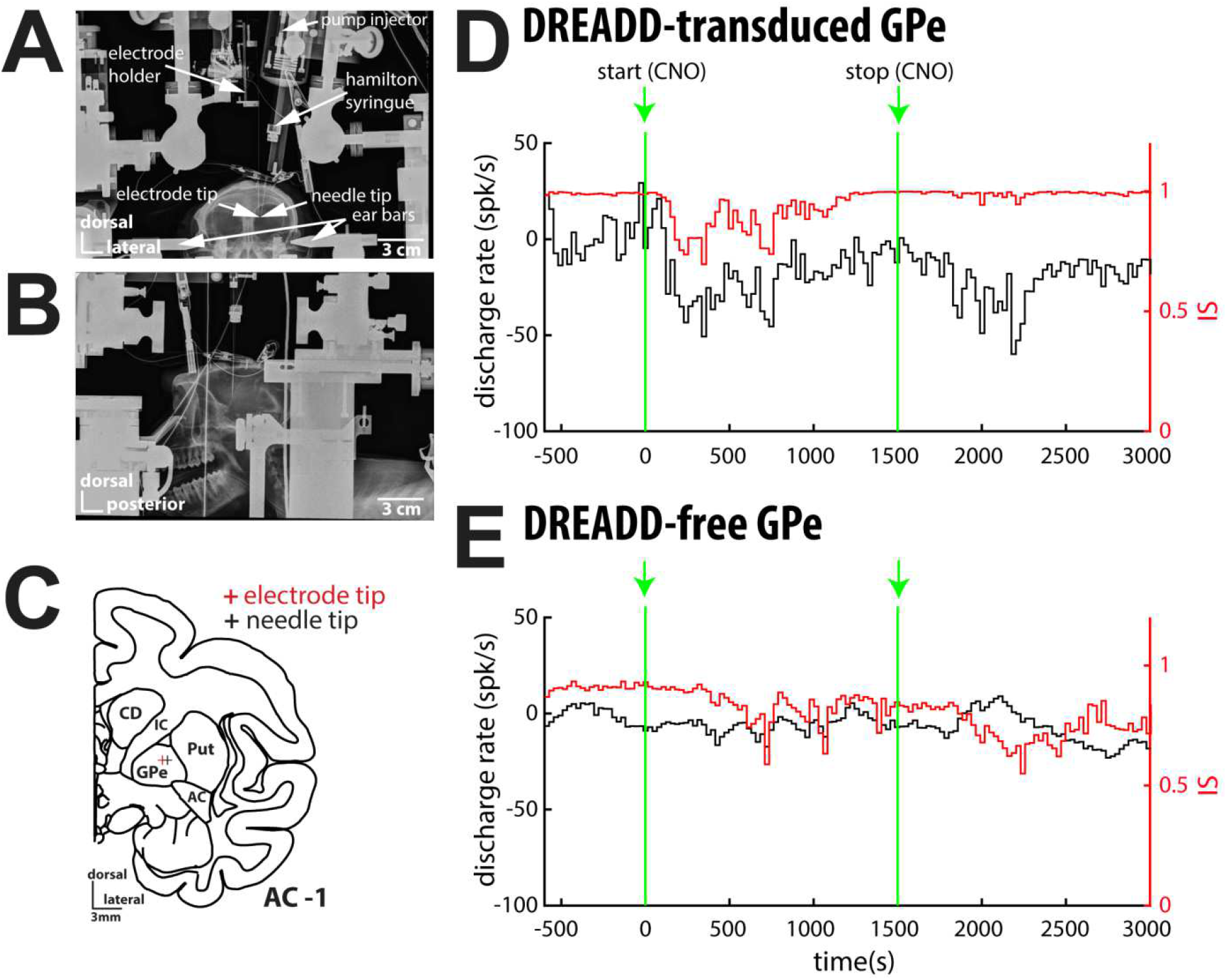
Recording configuration and examples. **(A)** Coronal and **(B)** sagittal X-ray images of the animal in the stereotaxic frame showing the apparatus during GPe unit recordings. **(C)** Representative coronal section - 1 mm from the anterior commissure (adapted from Martin & Bowden, 2000) indicating the location of both the electrode and needle tips in the left DREADD-transduced GPe during neuronal recordings. **(D and E**) perievent time histograms (PETHs) of two representative GPe units recorded following local CNO microinjection in DREADD-transduced **(D)** and DREADD-free **(E)**. Firing rate of each unit (left y-axis) is defined as the deviation from its baseline by subtracting the baseline firing rate (mean firing rate during the 10-min baseline period before CNO injection). Red traces indicate the evolution of the isolation score (IS, right y-axis) before, during and after CNO injection. 0 and 1500s correspond to CNO injection start and end (5μl CNO injection at 0.2 μl/min).

### Data analysis

Single-unit striatal activity was assessed by sorting spike trains from each 250-6000Hz band-pass filtered signal recorded within the GPe via offline spike detection and sorting method (Offline Sorter v4.4.2.0, Plexon Inc, Dallas, TX, USA). Threshold-crossing method for spike detection (negative voltage threshold trigger), principal component analysis for spike feature extraction and K-means clustering method (K-Means Scan with number of clusters between 1 and 5) were applied.

The isolation quality of each sorted unit before, during and after CNO injection was systematically graded by calculating its isolation score (IS, Joshua *et al.*, 2007) for each 30s-bin. IS ranges from 0 (i.e. highly noisy) to 1 (i.e. perfect isolation). Quantification of the isolation quality of the sorted units is extremely important. Indeed, poor quality isolation of the sorted units affects the calculated discharge rate (Lewicki, 1998; Joshua *et al.*, 2007). False negative errors, namely missing real spikes, decrease the calculated discharge rate. On the other hand, false positive errors, namely erroneously classifying noise events as spikes, lead to artificial increase in the discharge rate of the detected unit.

Only sorted units with mean IS ≥ 0.6 (well-isolated units) over their full recording span were included in the database. Moreover, we also identified and excluded the 30s-bins where IS ≥ 2 or ≤ −2SD of the mean IS (P < 0.05, empirical 68-95-99.7 rule) in order not to erroneously bias the results with high or low firing rate due to the poor isolation quality of a unit during particular bins.

Then, we calculated the perievent time histogram (PETH) for each well-isolated unit. Given that GPe neurons emit massive collaterals and their inhibitory GABAergic nature (Kita & Kitai, 1994; Bevan *et al.*, 1998; Sato *et al.*, 2000; Sadek *et al.*, 2007), one might expect that DREADD-based inhibition of the GPe neurons located in the vicinity of the recorded GPe will provoke some increases in its activity. A unit was therefore defined as a modulated unit when its activity during and/or after CNO injection was ≥ 2 or ≤ - 2 SD of the baseline firing rate (i.e., mean firing rate during the 10-min baseline period before CNO injection). Moreover, for each PETH, we also determined at each 30s-bin the polarity (increase vs. decrease) of the activity compared to its baseline and re-calculated the activity period (i.e., during and after CNO injection) when only considering the increases or decreases in activity. Unit was considered to significantly increase or decrease its activity when its activity period ≥ 2 or ≤ - 2 SD of the baseline firing rate, respectively.

### Histology and DREADD extent analysis

At the end of the experimental sessions, animals were deeply anesthetized with a lethal dose of pentobarbital (50 mg/kg) and transcardially perfused with 0.9% physiological saline solution. The brain was removed for each monkey and the two hemispheres were divided and cut into three parts each. These tissues were post fixed in a large volume of 4% buffered paraformaldehyde solution for one week at 4°C and cryoprotected in successive baths of 20% and 30% sucrose solution diluted in 0.1M phosphate-buffered saline (PBS) at 4°C until they sunk. Finally, brains were frozen by immersion in an isopentane bath at −55°C for 5 minutes and stored at −80°C. The entire basal ganglia (and more specifically the GPe) were cut in the coronal plane on a Leica 3050S cryostat into 50µm serial free-floating sections (24 series) collected in PBS-Azide 0.2% and store at 4°c until they were processed for mCherry immunoreactivity.

One series for each hemisphere was used for the staining. Briefly, endogenous peroxidases were first blocked by a peroxidase block reagent incubation of 10min (DAKO Peroxydase Block reagent, #S202386, Aglient, France). After 2 rinses with 0.1M PBS, sections were blocked in a solution of BSA 2%-TritonX100®0.3%-PBS 0.1M for 30 minutes before being incubated overnight in rabbit monoclonal anti-mCherry antibody (Abcam, ab213511; 1:1.000 dilution in BSA 0.2%-TritonX100®0.3%-PBS 0.1M). After rinsing sections with PBS, the staining was revealed by 30 min incubation in a HRP-anti rabbit polymer system (DAKO REAL Envision™ Kit, #K400311-2, Agilent, France) followed by a DAB revelation of few seconds (DAKO DAB Kit, #K346811-2, Agilent, France). Free-floating sections were mounted on gelatine-coated slides, counterstained with 0.2% cresyl-violet solution, dehydrated and coverslipped. Then, high-resolution whole slide color images were acquired with a Panoramic SCANNER (3D Histech, Hungary) at x20 magnification with 5 layers in extended picture. These high-resolution pictures were used to localize the injection and recording sites based on the Nissl stain which allowed the visualization of traces due to the passage of syringes and electrodes, and also to analyse DREADD expression.

The extent of mCherry staining within the GP (i.e., both the external and internal segments of the globus pallidus) was quantified with threshold surface analysis (Arotcarena *et al.*, 2019) using Mercator software (Mercator V7.13.4, Explora Nova, France). The mCherry fluorescent protein tag was fused to the hM4Di DREADDs; therefore the extent of mCherry staining reliably reflected the expression of hM4Di DREADDs. Briefly, GP was first delineated for each serial section of the series. The color thresholding tool of the software was used to select the threshold corresponding to the mCherry staining. The software extracted the surface corresponding to both the GP and the defined threshold, and the extent of mCherry staining was defined as the percentage of mCherry staining within the GP cross section surface. The same threshold surface analysis was performed for 2mm diameter circles centered on the presumed (visually identified on GP sections) viral construct injection sites.

### Software and Statistics

The data from the two monkeys were pooled since no significant difference was detected between them. All the data and statistical analyses were done using custom-made MATLAB R2016a routines (Mathworks, Natick, MA, USA). Wilcoxon signed rank test was used for statistical comparisons of paired groups. Statistics presented in this report, if not specified, use mean ± standard error to the mean (SEM) and the criterion for statistical significance was set at P < 0.05 for all statistical tests.

## Results

### Database

We recorded 19 and 8 well-isolated (IS ≥ 0.6) GPe units (monkeys F and E) following local intra-GPe microinjection (in the proximity of the electrode tip) of CNO in DREADD-transduced and DREADD-free GPe. Fig.1D and E depicts two representative examples of neurons showing the evolution of their firing rates following CNO injection in DREADD-transduced (Fig.1D) and DREADD-free (Fig.1E) GPe. As illustrated, a substantial decrease of the firing rate was observed following local intra-GPe microinjection of CNO in the DREADD-transduced GPe, while isolation quality of the unit remained stable over time (Fig.1D). In contrast, no consistent change in the firing rate was found following local intra-GPe microinjection of CNO in the DREADD-free GPe, while isolation quality of the unit remained stable over time (Fig.1E).

### Population averaged analysis

Fig.2A and B (upper left panels) depicts the mean PETHs for DREADD-transduced and DREADD-free GPe, respectively. The mean PETH is calculated as the arithmetic average of all the Z-normalized PETHs. For Z-normalization, we used the mean and SD firing rate of the 10-min baseline period before CNO injection. In DREADD-transduced GPe, we found 6/19 (31.6%) modulated units following CNO injection (3 during CNO injection and 3 after CNO injection, Fig.2A, upper middle panel), resulting in a significant (Wilcoxon signed rank test, Zval = 2.1557, P = 0.0311,) decrease in activity after CNO injection (−7.13 ± 1.68Hz, Fig.2A, upper right panel). In contrast, there was no modulated unit following CNO injection in DREADD-free GPe (Fig.2B, upper middle panel). Consistently, we did not find any significant net change in activity following CNO injection in DREADD-free GPe (Fig.2B, upper right panel).

**Fig.2.**
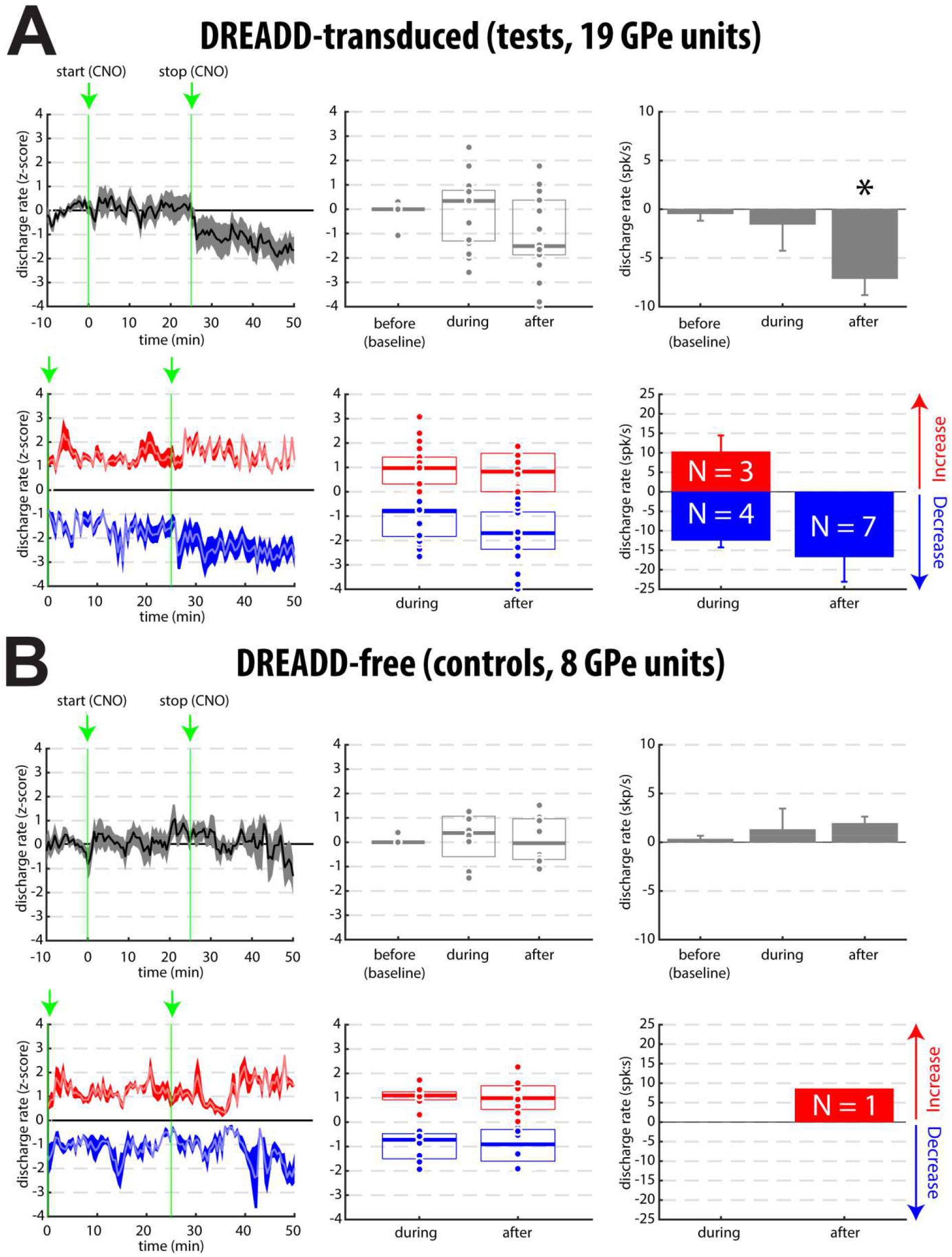
Population analysis. Effect of a local CNO microinjection on the pallidal activity in **(A)** DREADD-transduced and **(B)** DREADD-free GPe. **Left panels**, population responses subsequent to local CNO microinjection regardless **(top panels)** or not **(bottom panels)** of the polarity of the modulations (increases vs. decreases) in activity. Population response to CNO injection was defined as the average of all the Z-normalized PETHs. For each unit, the polarity (increase vs. decrease) of its activity at each 30s-bin was defined according to the baseline, time-varying increase/decrease balance is therefore depicted during and after CNO injection. 0 and 25s correspond to CNO injection start and end (5μl CNO injection at 0.2 μl/min). **Middle panels**, box plots showing the Z-normalized activity of each unit before, during and after CNO injection, regardless **(top panels)** or not **(bottom panels)** of the polarity of the modulations in activity. For each unit, activity period corresponds to the mean of the Z-normalized activity before, during and after CNO injection. In each box, the bold line indicates the median, and the bottom and top edges of the box indicate the 25th and 75th percentiles, respectively. **Right panels**, bar plots showing the mean deviation of the firing rate from the baseline regardless **(top panels)** or not **(bottom panels)** of the polarity of the modulations in activity. For each unit, the firing rate was defined as the deviation from its baseline by subtracting the baseline firing rate (mean firing rate during the 10-min baseline period before CNO injection) and averaged over the period (before, during and after CNO injection). Mean deviation of the firing rate was defined as the average of the deviation of the firing rate of all the units [top panels, * indicates significant difference (p < 0.05) compared to the baseline] or only the units with a significant increase or decrease in activity [bottom panels, N indicates the number of units significantly increasing or decreasing their activity]. Error bars represent SEMs

When examining time-varying increase/decrease balance of activity following CNO injection in the DREADD-transduced GPe (Fig.2A, lower left panels), we found that 3/19 (15.8%) and 4/19 (21.1%) units increased and decreased, respectively, their activity during CNO injection (Fig.2A, lower middle panels). In addition, 7/19 (36.8% - 4 of them also decreased their activity during CNO injection) units decreased their activity after CNO injection (Fig.2A, lower middle panel). Overall, 9/19 (47.4% - note that one unit increased and decreased its activity during CNO injection) units modulated their activity following CNO injection in DREADD-transduced GPe. These results suggest that neuronal activity of the well-isolated units exhibited diverse patterns in timing and polarity (increase/decrease) of firing rate modulations during and after CNO injection. Nevertheless, although increase and decrease levels were relatively similar during CNO injection (+10.27 ± 4.19Hz and –12.49 ± 1.78Hz, respectively), decreases in activity after CNO Injection, in addition to be exclusive, were also stronger (−16.75 ± 6.33Hz, Fig.2A, lower right panel). In contrast, only one unit increased its activity after CNO injection in DREADD-free GPe (+9.44Hz, Fig.2B, lower middle and right panels)

### DREADD expression

Extent of mCherry staining (based on the threshold surface analysis, see Materials and Methods) within GP of monkey F is represented in Fig. 3A. Threshold surface analysis revealed that mCherry staining surface in GP was larger around AC0, AC-1 and AC-2 (and maximal at AC-1, Fig. 3B). Also, when examining mCherry staining in the vicinity of the viral construct injection sites located in the GPe (2mm diameter circles centered on the viral construct injection sites), mCherry staining surface reached up to ~24% (Fig. 3C, left). Moreover, we observed a substantial number of mCherry-labeled GPe neurons (Fig. 3C, right). These results therefore indicate that hM4Di DREADDs fused to mCherry were expressed at high level in the GPe neurons located in the vicinity of the viral construct injection sites, which were also the recording sites.

**Fig.3.**
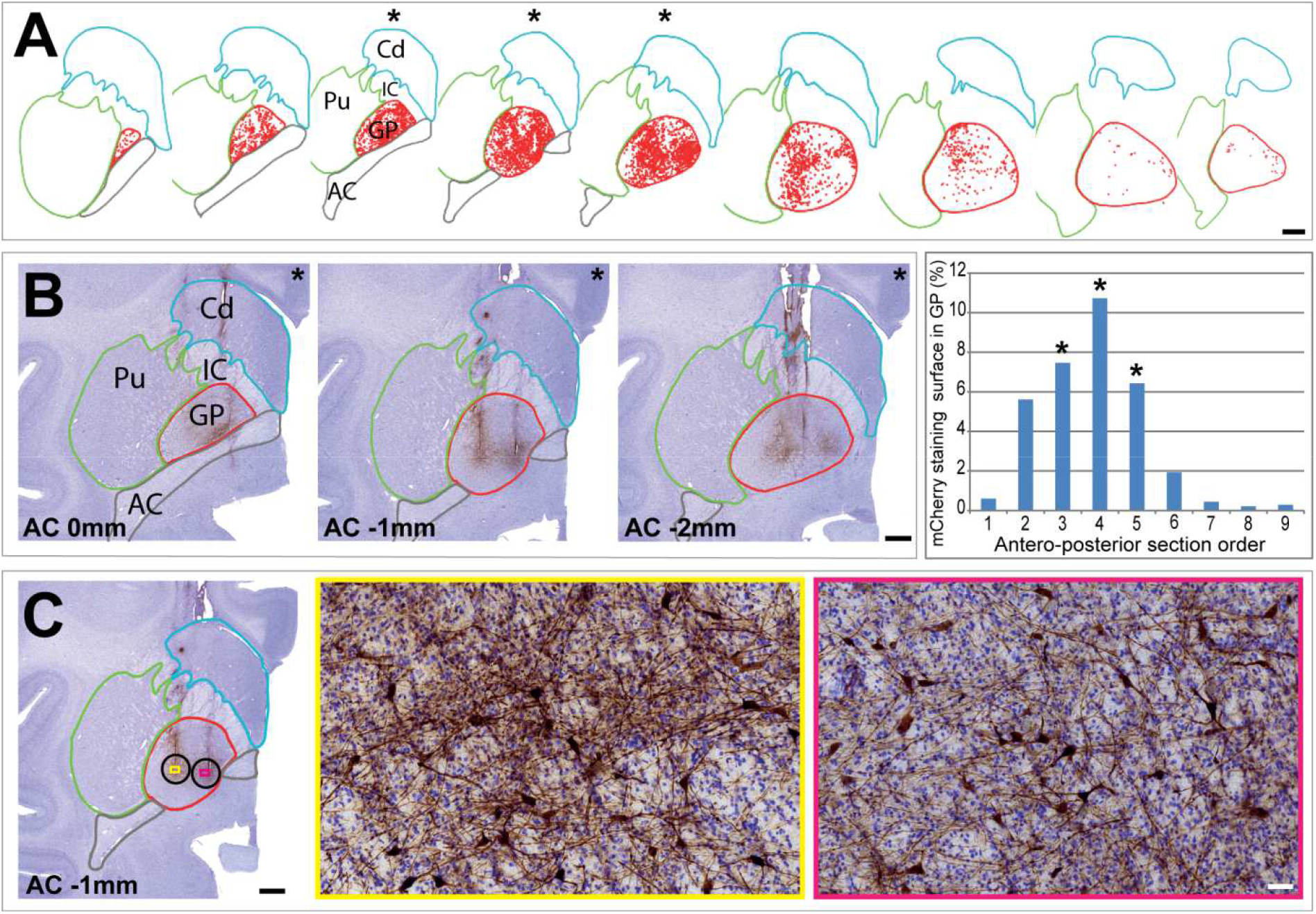
mCherry immunoreactivity. **(A)** Antero-posterior representation of the extent of mCherry staining within the GP (using threshold surface analysis, scale bar: 2000µm). **(B)** Illustration of mCherry staining patterns at AC0, AC-1 and AC-2 (left, scale bar: 2000µm) and quantification of mCherry staining surface for the different GP sections (right). Asterisks indicate the three sections with the highest mCherry staining surface in GP, namely AC0, AC-1 and AC-2**. (C)** Microscopic illustrations (yellow and pink boxes) of the density of mCherry-labeled GPe neurons located in the vicinity of two viral construct injection sites [2mm diameter circles (black) centered on the viral construct injection sites]. Scale bar: 2000µm (left) and 50µm (right). Abbreviations: AC, anterior commissure; Cd, caudate nucleus; IC, internal capsule; GP, globus pallidus; Pu, putamen.

## Discussion

This study is the first to demonstrate *in vivo* DREADD-based inhibition of the neuronal activity in the NHP (Fig., 1D and Fig.2A). Our results reinforce the view that DREADD technique can be used as a powerful tool to remotely, reversibly and repeatedly manipulate the neural mechanisms underlying higher brain functions, not only in rodents, but also in NHPs. Besides, given the unique value of NHP models in translational research, our results are more easily translatable to human. In neurosciences, traditional and available techniques in NHPs to examine the causal link between structure and function are based on lesion or pharmacological approaches which are highly invasive, if not irreversible. Development of the DREADD technique in NHP model is therefore fundamental for basic and translational neuroscience research.

Decreases in activity following local intra-cerebral CNO microinjection, albeit significant, were moderate (Fig., 1D and Fig.2A). In any case, we were not able to silence the activity of the neurons located in the DREADD-transduced GPe. Our post-mortem histochemical analysis revealed that hM4Di DREADDs were expressed at high level in the GPe neurons located in the vicinity of the viral construct injection and recording sites (Fig. 3). However, our approach did not allow us to examine the subcellular localization of the DREADDS (i.e., intracellular space vs. plasma membrane). In a recent NHP study, Galvan and colleagues demonstrated that the pattern of subcellular expression of DREADDS in monkeys depends on the DREADD/tag combination (Galvan *et al.*, 2019). In particular, they showed that using hM4Di DREADD fused to mCherry (as we did here) leads to a low proportion of hM4Di at the plasma membrane of the monkey, but not the rodent, neurons. In contrast, using the same hM4Di DREADD fused to the HA-tag leads to a robust plasma membrane expression in both monkeys and rodents neurons. Further studies should be therefore conducted to test the efficacy of different DREADD/tag combinations for silencing the neuronal activity. However, since behavioral effect has already been observed in monkeys using the hM4Di-mCherry combination (Grayson *et al.*, 2016; Upright *et al.*, 2018), it is possible that a small fraction of hM4Di at the plasma membrane is sufficient for decreasing the activity of DREADD-transduced neurons. Finally, although we used high (mM) concentration of CNO for microinjections that might lead to off-target effects (Gomez *et al.*, 2017), we found no robust modulations of activity in the DREADD-free GPe. Nevertheless, one can assume that higher concentrations will produce stronger decreases in activity in the DREADD-transduced GPe.

Surprisingly, we found that neurons in DREADD-transduced GPe exhibited diverse timing and directional (increase or decrease) patterns of discharge modulations following local CNO microinjection (Fig.2A, lower panels). GPe neurons possess local axon collaterals (Kita & Kitai, 1994; Bevan *et al.*, 1998; Sato *et al.*, 2000; Sadek *et al.*, 2007). Given the GABAergic nature of the GPe neurons, it is possible that inhibition of DREADDs-transduced GPe neurons in the vicinity of the recorded GPe neurons leads to increases in the activity of the recorded neurons or at least to the cancelation/reduction of the expected decrease in activity due to CNO exposure. Nevertheless, we demonstrated that decreases in activity prevailed in the DREADD-transduced GPe after CNO microinjection (Fig.2A, lower panels), thus indicating DREADD-based inhibition of NHP pallidal neurons.

The ultimate benefit of DREADDs technique is to remotely, reversibly and repeatedly manipulate (with minimal invasiveness) the neuronal activity via systemic injection of specific ligand. Here, our goal was to unambiguously show (using a straightforward strategy) that DREADDs can reduce firing rate of NHP neurons when activated by a specific ligand. When systemically injected, the use of CNO is problematic. Indeed, CNO hardly crosses the blood-brain barrier (Gomez *et al.*, 2017; Raper *et al.*, 2017) and can be *in vivo* converted to clozapine (Jann *et al.*, 1994; Chang *et al.*, 1998; MacLaren *et al.*, 2016; Gomez *et al.*, 2017; Raper *et al.*, 2017) which shows high blood-brain permeability and high affinity to muscarinic-based DREADDs (Ji *et al.*, 2016; Gomez *et al.*, 2017). However, clozapine is capable of binding to a variety of endogenous receptors (Galvan *et al.*, 2018). This is why we favored local intra-cerebral CNO microinjections to make sure our results are not confounded by endogenous non-DREADD target actions of the clozapine. Recently, new DREADD ligands (i.e., JHU37152 and JHU37160) with excellent brain penetrance and high *in vivo* DREADD potency (including in NHP) has been developed (Bonaventura *et al.*, 2019). To close the loop, further experiments should be therefore conducted to ensure that systemic injections of these effective DREADD ligands (Bonaventura *et al.*, 2019) could also inhibit NHP neuronal activity.

## Acknowledgements

This study was supported by the French National Research Agency (ANR) and the French National Center for Scientific Research (CNRS) to M.D and T.B. We thank Dr. Gilad Silberberg and Dr. Liliya Iskhakova for careful reading and comments. We are grateful to Dr. Nicolas Mallet and Dr. Arthur Leblois for insightful scientific discussions.

## Competing Interests

The authors declare no conflict of interest.

## Data Accessibility

The data are available from the corresponding author upon request.

## Author Contributions

M.D., E.B., T.B. designed the research; M.D., T.H.N., H.O., N.B., S.D. performed the research; M.D., analyzed the data; M.D., T.B. wrote the paper; All authors read and approved the final version.

## Abbreviations

AAV: adeno-associated virus
CNO: clozapine-N-oxide
DREADDs: Designer Receptors Exclusively Activated by Designer Drugs
GPe: external globus pallidus
IM: intramuscular
IS: isolation score
NHP: non-human primate
PETH: perievent histogram
PDS: phosphate-buffered saline
SD: standard deviation
SEM: standard error to the mean

